# Structural insights into human TFIIIC promoter recognition

**DOI:** 10.1101/2023.05.16.540835

**Authors:** Wolfram Seifert-Davila, Mathias Girbig, Luis Hauptmann, Thomas Hoffmann, Sebastian Eustermann, Christoph W. Müller

**Author notes:** These authors contributed equally (Wolfram Seifert-Davila, Mathias Girbig).

## Abstract

Transcription factor IIIC (TFIIIC) recruits RNA polymerase (Pol) III to most of its target genes. Recognition of intragenic A- and B-box motifs in tRNA genes by TFIIIC modules τA and τB is the first critical step for tRNA synthesis but is mechanistically poorly understood. Here, we report cryo-EM structures of the human 624 kDa TFIIIC complex unbound and bound to a tRNA gene. The τB module recognizes the B-box via DNA shape and sequence readout through the assembly of multiple winged-helix domains. TFIIIC220 forms an integral part of both τA and τB connecting the two subcomplexes via a ∼550 amino acid residue flexible linker. Our data provide a structural mechanism by which high-affinity B-box recognition anchors TFIIIC to promoter DNA and permits scanning for low-affinity A-boxes and TFIIIB for Pol III activation.

## INTRODUCTION

Nuclear genes are transcribed by DNA-dependent RNA polymerases, which are recruited to promoter sequences by general transcription factors (GTF)^1^. Transcription factor IIIC (TFIIIC) – a six-subunit protein complex with a molecular weight (MW) of 624 kDa in humans – facilitates transcription of type 1 (5S rRNA) and type 2 (e.g. tRNAs) RNA polymerase (Pol III) target genes^2–5^. Whereas type 1 genes also require TFIIIA, TFIIIC is the first GTF that binds type 2 genes and subsequently recruits the three-subunit TFIIIB complex, composed of BRF1, BDP1 and TBP. TFIIIB binds upstream of the transcription start site and aids Pol III to bind and open the promoter DNA^6^. TFIIIC is composed of two subcomplexes, τA and τB, that bind two gene internal DNA motifs: A-box and B-box^7, 8^. τA binds the A-box with low affinity and is required to recruit TFIIIB to the promoter DNA^6, 9^. τB interacts with the B-box with high affinity and, therefore, is essential for target DNA recognition of type 2 genes^9^.

Deregulation of the Pol III transcription apparatus was reported in various cancers (reviewed in Ref.^10^), and increased expression- and protein levels of TFIIIC subunits were found in ovarian cancers^11^ and during tumoral transformation^12^. Independently of its role in recruiting Pol III to its target genes, TFIIIC recognizes ‘extra TFIIIC’ (ETC) sites in various eukaryotes^13–15^ and functions in genome organization by acting as a chromatin insulator^16, 17^ and via interaction with the architectural proteins cohesin and condensin^18–21^. TFIIIC also possesses histone acetylation activity and, together with the insulator protein CTCF, was shown to mediate long-range chromatin looping^22, 23^.

TFIIIC must possess a high degree of structural elasticity since A- and B-box binding implies spanning the entire tRNA gene body^24, 25^, whose size varies from 70-108 basepairs (bps) in *Homo sapiens* and 71-133 bps *in Saccharomyces cerevisiae*. Owing to this structural elasticity, our molecular understanding of how TFIIIC is organized has been limited to structures of subassemblies of τA and τB from yeast^26–30^. Hence, it remained enigmatic how τA and τB interact with each other and, thereby, form a stable and, at the same time, flexible molecular assembly. Structural insights into how TFIIIC recognizes its target genes were missing, and no structures of human TFIIIC or its building blocks have been experimentally determined.

## RESULTS

### Structure determination of human TFIIIC

Human TFIIIC consists of τA subunits TFIIIC35, TFIIIC63 and TFIIIC102 and τB subunits TFIIIC90, TFIIIC110 and TFIIIC220 (Figure 1A). To obtain structural insights into human TFIIIC, we co-expressed its subunits in insect cells and confirmed by mass photometry (MP) that the purified sample corresponds to the expected size of the intact human TFIIIC complex (MW_observed_ = 654 ± 52 kDa, MW_expected_ = 624 kDa) (Figure S1A-S1C). Reconstitution with a type II promoter DNA encoding the human TRR-TCT3-2 tRNA (tRNA_Arg_) gene (MW = 61 kDa) gave rise to a 701 ± 46 kDa complex indicating cognate DNA binding activity of the complex at low nanomolar concentrations (Figure S1D).

**Figure 1.**
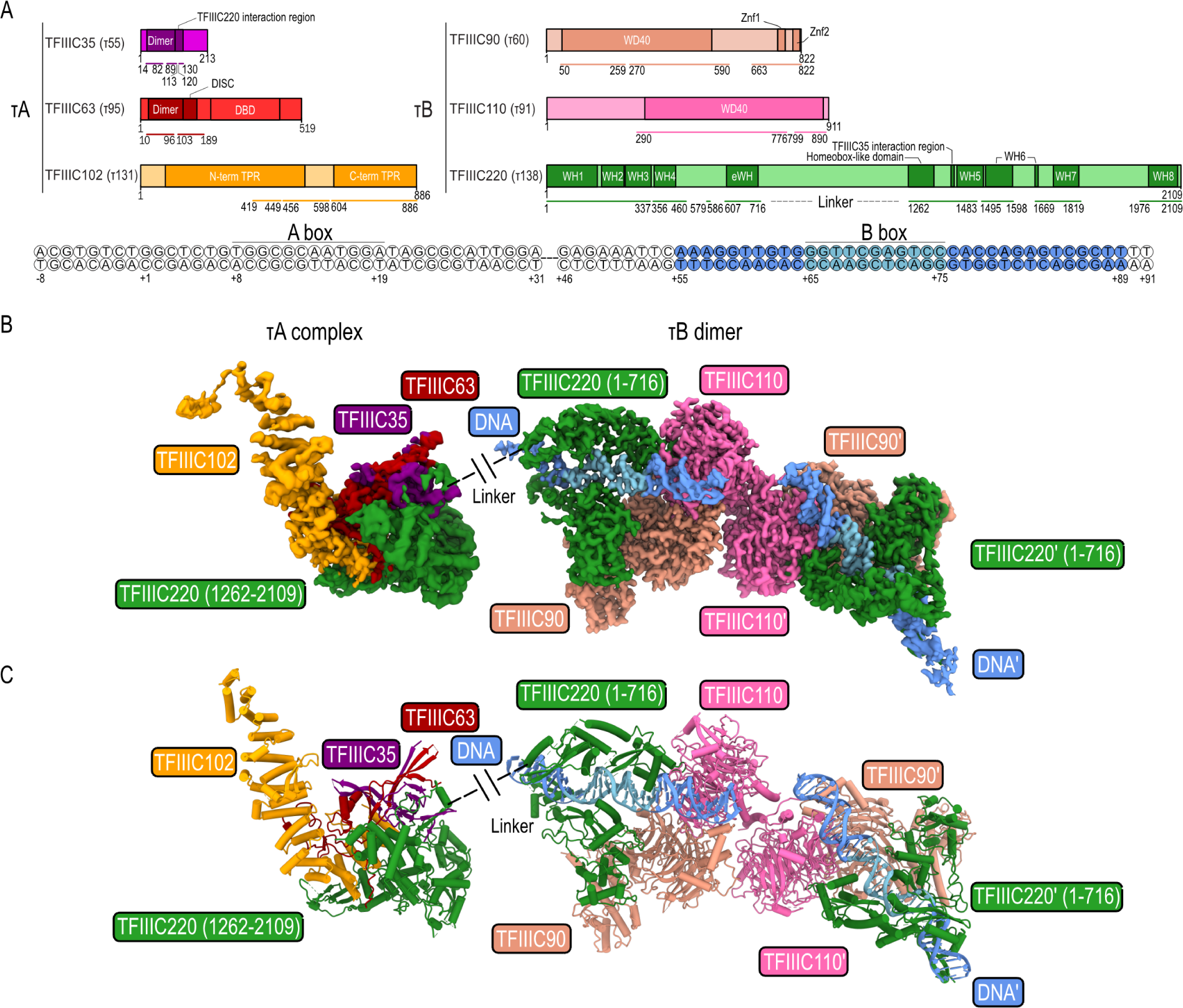
Cryo-EM structure of human TFIIIC. **(A)** Domain diagram of human TFIIIC subunits (top) and the tDNA used for cryo-EM analysis (bottom). Yeast homologs are given in parentheses. Colored bars: built regions, colored circles: modeled DNA. DBD - DNA binding domain, TPR - tetratricopeptide domain, WD40 - WD40 repeat domain, WH - winged-helix domain, eWH - extended winged-helix domain. **(B)** Cryo-EM maps of human τA and dimeric, DNA-bound τB. **(C)** Atomic models of human τA and DNA-bound τB.

We determined cryo-EM structures of human TFIIIC alone and in complex with the TRR-TCT3-2 tRNA gene. In both cases, we obtained two distinct sets of particles for human τA and τB. Separate cryo-EM reconstructions at 3.8 and 3.4 Å resolution (DNA-unbound) and at 3.5 and 3.2 Å resolution (DNA-bound) (Figure S2) enabled us to build and refine atomic models of human τA and τB, respectively (Figure 1B and 1C; Figure S3A-S3D; Table S1).

To our surprise, we observed a dimeric structure of τB in both the DNA-unbound and DNA-bound 3D reconstructions (Figure 1B and S4A). The two τB subcomplexes are held together by apolar and polar interactions between two copies of subunit TFIIIC110 (Figure S4B). TFIIIC dimerization is likely to be concentration dependent since our MP measurements did not show clear indications for dimer formation at nanomolar concentrations. The observation that TFIIIC dimerizes under cryo-EM conditions should be treated with care because we were unable to evaluate TFIIIC110 dimerization in solution due to a limited amount of protein sample. Since TFIIIC also functions in 3D genome organization^22, 23^, one could, nevertheless, speculate that DNA-bound τB dimerizes at tRNA gene clusters and ETC sites where it may help to stabilize long-range chromatin loops. For the following molecular description of human τB, we will, however, only focus on the τB monomer.

### Human τB core structure

The core of human τB is formed by TFIIIC90 and TFIIIC110 (τ60 and τ90 in yeast), which stably associate with each other via their WD40 domains (Figure S5A). We could also assign the N-terminal third (1-716) of subunit TFIIIC220 (τ138 in yeast) featuring four winged-helix (WH) domains and one extended WH (eWH) domain to human τB (Figure 1B and 2). TFIIIC220-WH1 (1-163) binds two zinc-fingers of the TFIIIC90 C-terminal domain (Figure S5B) and engages with the WH2 domain (175-247). On the other edge of τB, the TFIIIC110-WD40 domain stabilizes the TFIIIC220-WH4 (367-437) domain.

**Figure 2.**
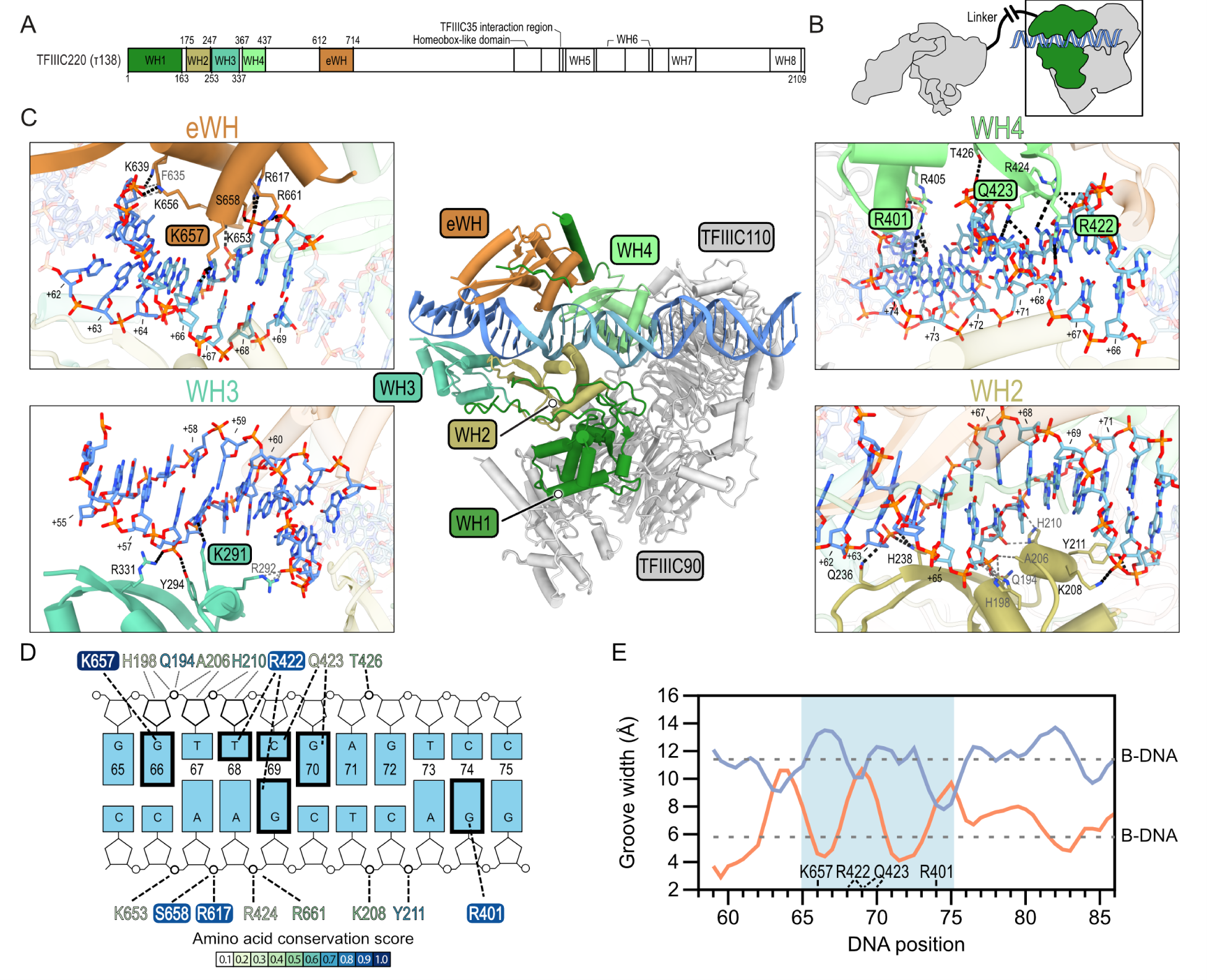
Interaction of τB with B-box promoter DNA. **(A)** Domain architecture of TFIIIC220. N-terminal, DNA-binding domains are colored. **(B)** Schematic of hTFIIIC bound to DNA. Green: TFIIIC220 N-terminal moiety. **(C)** Centre: atomic model of human τB bound to DNA (light blue: B-box). Contacts between WH2, WH3, WH4 and eWH with the DNA are shown as close-up views. Black dashed lines: hydrogen (H)-bonds, grey dashed lines: apolar contacts. H-bond forming residues are labeled and amino acids that bind the DNA-bases are highlighted in colored boxes. **(D)** Nucleotide plot of the B-box DNA-TFIIIC220 interaction. H-bonds are depicted as bold, dashed lines and apolar contacts are shown as grey, thin lines. Bases that are contacted directly are highlighted as bold frames. Contacting residues are colored according to their Scorecons^78^ conservations scores. Residues with a conservation score > 0.9 are framed. **(E)** DNA groove width analysis of the TFIIIC220-bound tDNA, computed with the Curves+ software^79^. The B-box motif is highlighted in light blue. Orange and blue curves show DNA-minor- and major groove widths, respectively. Dotted lines represent minor- and major groove widths of ideal B-DNA. TFIIIC220 residues that form base-specific contacts are depicted above the nucleotide positions.

Hence, TFIIIC90-TFIIIC110 form – with the help of the TFIIIC220 WH1 domain and the C-terminal zinc-fingers in TFIIIC90 – a stable scaffold, which stabilizes the WH2 and WH4 domains. These two domains were, therefore, already visible in the absence of DNA, whereas the WH3 and eWH domains could not be visualized in the DNA-unbound human τB cryo-EM map (Figure S4A).

### Structural basis of B-box DNA recognition

The cryo-EM structure of DNA-bound human τB enabled us to unveil the molecular basis of B-box DNA recognition. The 3.2 Å cryo-EM map of DNA-bound τB was of sufficient quality to assign 35 bps (55-89) of the DNA sequence (Figure 1A). Upon DNA binding, the TFIIIC220-WH3 (253-337) and the eWH-domains (612-714) become ordered and facilitate B-box (65-75) DNA recognition together with WH2 and WH4 (Figure 2 and Figure S6A). The DNA is, downstream of the B-box, additionally stabilized by a positively-charged surface of TFIIIC110 (Figure S5C). The lack of this subunit in human has been associated with a transcriptionally inactive TFIIIC form^31^. In yeast, the subunit τ91 (TFIIIC110 in human) has been shown to cooperate with τ138 (TFIIIC220 in human) for DNA binding^32^. The WH domains huddle against the DNA from both sides and form several apolar and polar contacts with the phosphate backbone and bases of the B-box DNA (Figure 2C and S6A). By analyzing the B-box DNA sequence conservation, we found that the DNA bases that are contacted in a base-specific manner (G66, T68, C69, G70, and G74) are not only conserved in human tDNAs but also across a wide range of eukaryotes (Figure S6B). In TFIIIC220, the basic amino acids (R401, R422, K657) that form base-specific contacts with the B-box DNA are also highly conserved (Figure 2D). Notably, biochemical DNA-binding studies with yeast TFIIIC and the SUP4 gene have shown that the mutations C56→G and G57→C (corresponding to C69 and G70 in the human TRR-TCT3-2 tRNA gene) caused a 370- and 56-fold decrease, respectively, in DNA-binding affinity^33^. Our structural analysis, together with these previously conducted experiments, thus, confirm the importance of the recognition of these conserved bases. Interestingly, the geometry of the DNA is also substantially altered nearby the bases that appear to be contacted in a conserved manner (Figure 2E). K657 (eWH) binding induces the opening of the DNA major groove and compaction of the corresponding minor groove whereas R422/Q423 and R401 (WH3) widen the DNA minor groove up to ∼11 Å and 10 Å, respectively. Hence, TFIIIC initiates recognition of the tDNA promoter in an evolutionary conserved manner via a combined DNA shape and sequence readout of the B-box DNA motif.

### TFIIIC220 connects τA and τB with a flexible linker

Our cryo-EM analysis yielded a high-resolution structure of human τA, which, however, remained in the DNA-unbound state in the presence of the tRNA gene (Figure 3). This observation is in agreement with biochemical studies on human TFIIIC, where the TFIIIC2 component composed of 5 subunits with molecular weights of 230, 110, 102, 90 and 63 kDa^31, 34, 35^ interacts exclusively with B-box and is deficient in interacting with A-box^36–38^. However, upon the addition of the TFIIIC1 component, which full composition remains unclear, but possesses human BDP1^39^, the protection against DNase I expands to the A-Box and the transcription start site. On the other hand, in yeast TFIIIC, it has been shown that τA binds the DNA with low affinity and in an unspecific manner^9, 27, 40^.

**Figure 3.**
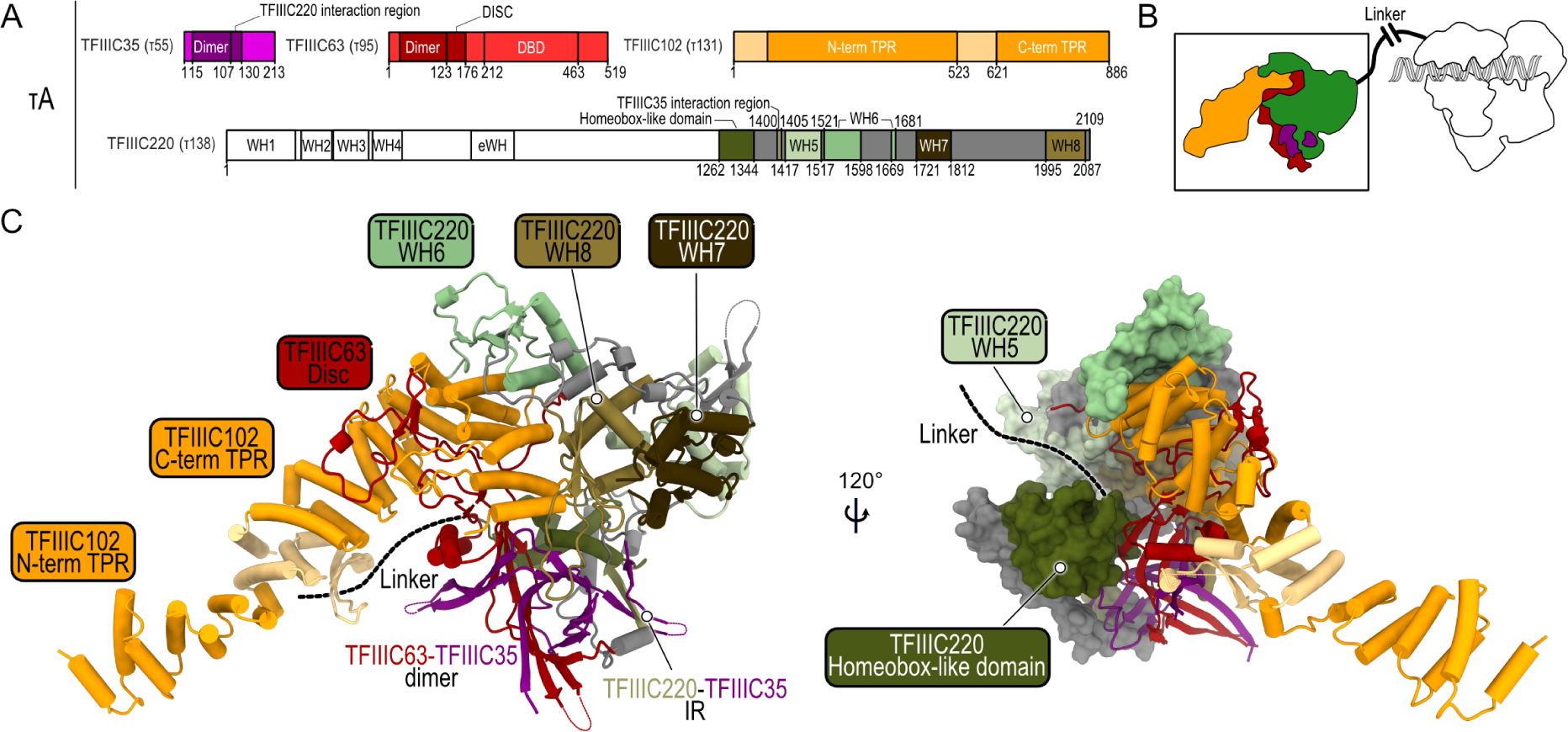
The C-terminal region of TFIIIC220 is an integral part of the τA subcomplex. **(A)** Diagram of the protein domains (colored) that form a stable τA subcomplex. For TFIII220: unmodelled regions are colored grey. **(B)** Schematic representation of τA (colored) and τB (white) connected by a flexible linker. **(C)** Atomic model of τA showing the C-terminus of TFIIIC220 as cartoon (left) and as surface (right). The flexible linker connecting τA and τB is shown as black dashed lines.

When inspecting the 3D reconstruction of human τA, we found that 50% of the cryo-EM density corresponds to the τA subunits TFIIIC35, TFIIIC63, and TFIIIC102 (τ55, τ95, and τ131 in yeast) (Figure 3A). TFIIIC35-TFIIIC63 form a heterodimer that binds the C-terminal TPR domain of TFIIIC102 (Figure 3C). Human τA, thus, partly resembles its yeast counterpart but it lacks the phosphatase domain of τ55 (absent in human TFIIIC35) and the cryo-EM density for the TFIIIC63-DNA binding domain (DBD). To our surprise, the remaining ∼50% of the cryo-EM density belongs to the C-terminal half of the τB-subunit TFIIIC220 (Figure 3). The corresponding atomic model features a homeobox-like domain, interacting with the TFIIIC35-TFIIIC63 heterodimer, and four WH domains (WH5-WH8) (Figure S7), of which WH6 and WH8 directly contact TFIIIC102. TFIIIC220, thereby, partly occupies the same binding interface of TFIIIC102 as the TFIIIC63-DBD in yeast (Figure S8).

The C-terminal half of TFIIIC220 forms an integral part of human τA, suggesting that TFIIIC220 links τA and τB with a ∼550 residue-long linker between its N- and C-termini. To address this hypothesis, we used our cryo-EM data to measure the distance between those particles that contribute initially to the τA and τB reconstructions (see methods). Our measurement revealed a distinct distribution of experimental particle distances, peaking at 200 Å, which differs from a set of simulated, randomly distributed particles (Figure S9A and S9B), indicating that the τA particles are stably, and, at the same time, flexibly linked to the tDNA-bound τB fraction. This flexibility allows TFIIIC to bridge the variable distances between A-box and B-box present in different tRNA genes in the human genome (Figure S9C).

## DISCUSSION

Based on our structural analysis, we propose a multi-step mechanism of tRNA-gene promoter recognition via TFIIIC (Figure 4). The first step involves the high-affinity DNA sequence and shape readout of the B-box motif via an assembly cascade of WH domains. WH2 and WH4 of TFIIIC220 are an integral part of the rigid τB-core, formed by TFIIIC90-TFIIIC110 and the TFIIIC220-WH1 domain, and their conformation does not change upon tDNA-binding suggesting that they provide the platform for initial DNA recognition. We speculate that τB-WH2-WH4 mediates scanning of the tRNA-gene for the correct B-box motif. Additionally, the TFIIIC-tDNA interaction is further stabilized by the WD40 domain in TFIIIC110, which binds the tDNA downstream of the B-box via a positively charged groove. Subsequently, the WH3 and eWH domains recognize the upstream half of the B-box, and the WH domains, collectively, enclose the B-box motif and its upstream region.

**Figure 4.**
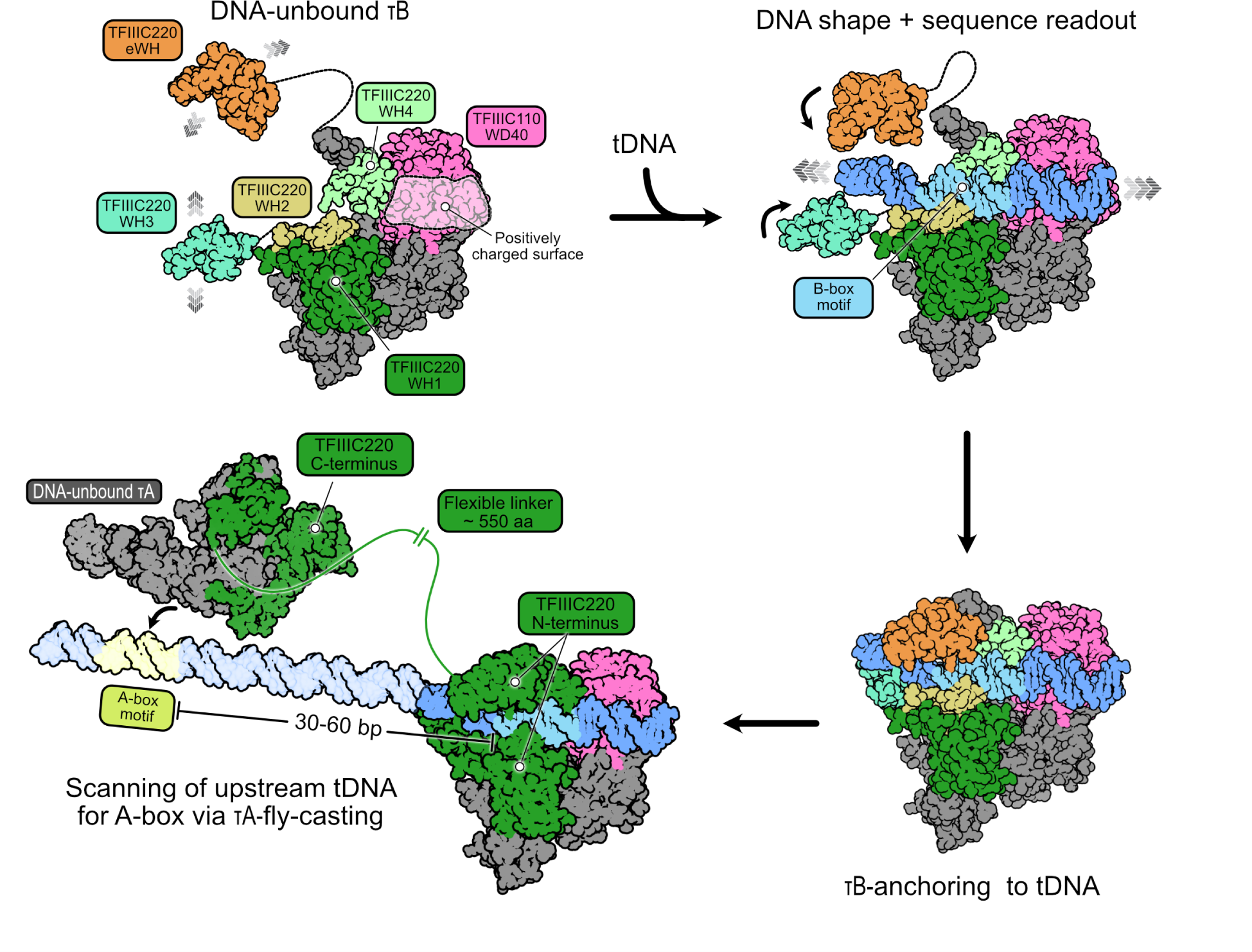
Model of TFIIIC promoter recognition. TFIIIC110 and TFIIIC90 along with the TFIIIC220-WH1, WH2, and WH4 domains form a rigid τB scaffold, while WH3 and eWH domains remain flexible. The DNA is initially recognized by the WH2 and WH4 domains, and subsequently by the WH3 and eWH domains via shape and sequence readout. Additionally, the TFIIIC110-WD40 positively charged pocket (transparent-white surface) helps to stabilize the interaction with the DNA molecule. Once TFIIIC is anchored to the tRNA gene, the τA subcomplex, connected by a ∼550 amino acid (aa) flexible linker to τB, will search for the A-box motif, separated from the B-box with a variable distance between 30-60 bp, by a fly-casting mechanism.

The concerted engagement of the B-box motif by τB WH domains at up- and downstream sites anchors TFIIIC to the tRNA gene and allows, via the flexible TFIIIC220 linker of 550 residues connecting τA and τB, subsequent scanning of the tRNA gene for low-affinity A-box motifs through a fly-casting mechanism. Composed of, in total, 9 WH domains, TFIIIC220 fulfils multiple key roles for TFIIIC-mediated promoter recognition. Whereas the N-terminal WH1-4 and eWH domains either directly or indirectly (WH1) mediate B-box recognition, the C-terminal WH5-WH8 domains form an integral part of τA and, thereby, serve as a hook that tethers τA and τB. The flexible TFIIIC220 linker between τA and τB also equips TFIIIC with the required elasticity to adapt to the variable distances between the A- and B-box motifs found across different tRNA genes.

It remains to be determined, how TFIIIC recognizes the A-box motif and recruits TFIIIB. TFIIIC functions as an assembly factor for TFIIIB^6^ but TFIIIB, vice versa, enhances the protection of the A-box DNA region in DNase I footprint experiments in yeast^41^, suggesting that TFIIIB and τA cooperatively recognize the promoter DNA. The absence of TFIIIB might explain why human τA remained in the DNA-unbound state in our 3D reconstruction. Yeast TFIIIB and TFIIIC interact with each other in solution via the TFIIIB subunit BRF1 and the τA-subunit τ131 (TFIIIC110 in humans)^30, 42–47^. It might, thus, be plausible that B-box-mediated anchoring of TFIIIC also allows τA to ‘fish’ for BRF1 in solution, thereby promoting TFIIIB assembly upstream of the transcription start site, which could, in turn, stabilize the association between τA and the A-box.

## Acknowledgments

We acknowledge support by K. Lapouge (EMBL Protein Expression and Purification Core Facility), M. Rettel (EMBL Proteomics Core Facility) and F. Weis and W. Hagen (EMBL Cryo-EM Service Platform). We also thank F. Baudin, H. Grötsch, H.K.H. Fung, and J. Weidenhausen for their support and discussion. WSD acknowledges support from the EMBL International PhD program. All authors acknowledge support by EMBL.

## Author contributions

WSD cloned and purified the hTFIIIC complex, collected and processed cryo-EM data, built the model and made figures. MG built and interpreted structural models and analyzed sequence conservation and DNA geometry. LH and TH helped with data processing, and TH implemented the required software. SE advised on cryo-EM sample preparation and supervised data processing. CWM initiated and supervised the project. WSD and MG wrote the manuscript with help of the other authors.

## Data and materials availability

Cryo-EM maps of human TFIIIC (map A to map E) have been deposited to the Electron Microscopy Data Base (EMDB) under accession codes EMD-XXXX. Atomic models of DNA-unbound human TFIIIC and human TFIIIC bound to DNA have been deposited to the Protein Data Bank under accession codes XXXX. Plasmids are available upon request and will be available without restrictions.

## METHOD DETAILS

### Protein expression

The six human TFIIIC-coding genes, codon-optimized for expression in insect cells, were ordered from DNASU. Two genes were edited via PCR to match the protein sequences found on UniProt^48^. The original GTF3C2 gene encoded 12 additional amino acids and GTF3C3 encoded a serine in position 70 instead of asparagine. Additionally, GTF3C6 encoded a serine in position 1, which was replaced by methionine. A restriction-free cloning protocol was applied to clone all the TFIIIC genes into a pACEBac vector. Using an overlap extension PCR cloning, a TEV-cleavable His-tag was added to the N-terminus of GTF3C2 and GTC3C5, followed by the addition of a 3XFLAG-tag to the C-terminus of GTF3C1. A six-gene TFIIIC expression plasmid was cloned via biGBac^49^. The genes belonging to τA and τB were cloned into the pBIG1a and pBIG1b plasmids, respectively, and the subcomplex-expression cassettes were cloned into pBIG2ab. A recombinant baculovirus was constructed following standard procedures.

Sf21 insect cells were used to express the TFIIIC complex. Sf21 cells were diluted to 0.5×10^6^ cells/ml and immediately infected with the TFIIIC-containing baculovirus. Sf21 cells were grown up to 72 h at 27 ⁰C or until their cell viability reached 90-95%. Cells were harvested by centrifugation at 700 x g, followed by a washing step with PBS, flash frozen and stored at −80 °C until use.

### Protein purification

Frozen pellets were resuspended for 2 h at 4 ⁰C while stirring gently in 3x lysis buffer (20 mM HEPES pH 7.5, 0.25 mM DTT, 500 mM NaCl, 2 mM MgCl2, 0.1 % NP-40, 10% glycerol, 0.25 mM DTT), supplemented with 1:1000 DNase I, SigmaFast EDTA-free protease inhibitors tablet per 20 g of cell pellet, 4 µl benzonase per 50 ml of lysis buffer. A lysis step to the resuspended pellet using a sonicator for 3 min was applied. The lysate was centrifuged at 30,000 × g for 1 h at 4 ⁰C to remove cell debris. Anti-FLAG M2 agarose beads were incubated with the remaining supernatant for 2 h on a rolling plate at 4 ⁰C. Beads were loaded onto an Econo-Column Chromatography column and washed with 30x column volumes (CV) of lysis buffer, followed by 25 CV of wash buffer 1 (20 mM HEPES pH 7.5, 500 mM NaCl, 10% glycerol, 0.1% NP-40, 0.25 mM DTT). The washed resin was incubated with 2x CV of elution buffer 1 (20 mM HEPES pH 7.5, 500 mM NaCl, 10% glycerol, 0.1 % NP-40, 0.25 mM DTT, 0.2 mg/ml FLAG peptide) for 30 min at 4 ⁰C. Wash buffer 2 (20 mM HEPES pH 7.5, 100 mM NaCl, 10% glycerol, 0.1 % NP-40, 0.25 mM DTT) was added to the eluted sample to dilute the initial salt concentration to 200 mM NaCl, then applied to a Capto HiRes Q 5/50 column pre-equilibrated in buffer A (20 mM HEPES pH 7.5, 200 mM NaCl, 5% glycerol, 5 mM DTT). Sample elution was carried out by applying a linear gradient from 0 to 100% buffer B (20 mM HEPES pH 7.5, 1 M NaCl, 5% glycerol, 5 mM DTT) over 70 ml. Peak fractions were analysed by SDS-PAGE. The fractions containing TFIIIC were pooled, concentrated up to 2mg/ml, buffer exchanged using storage buffer (20 mM HEPES pH 7.5, 200 mM NaCl, 5 mM DTT), flash frozen and stored at −80 °C until use or directly used for cryo-EM sample preparation.

### DNA oligonucleotide preparation

Only the template sequence is shown for simplicity. The Human TRR-TCT3-2 (tRNA_Arg_) gene (*ACGTGTCT*GGCTCTG<u>TGGCGCAATGGA</u>TAGCGCATTGGACTTCTAGATAGTTAGAGA AATTCAAAGGTTGTG**GGTTCGAGTCC**CACCAGAGTCG*CTTTT*, the A-box is underlined and the B-box is printed in bold, sequences flanking the Human TRR-TCT3-2 gene are in italics) was ordered from Sigma Aldrich. Non-template and template strands were annealed in H_2_O followed by heating to 95 ⁰C for 5 min before cooling down to 20 ⁰C at a rate of 1 ⁰C/min. The DNA mixture was applied to a Superdex 200 increase 3.2/300 column pre-equilibrated in DNA buffer (20 mM HEPES pH 7.5, 150 mM NaCl, 5 mM MgCl_2_, 5 mM DTT) to remove incomplete products and impurities. The fractions corresponding to the major peak were pooled and quantified by nanodrop. The sample was stored at −20 ⁰C until use.

### Mass-photometry sample preparation, data acquisition, and processing

High-precision microscope coverslips (24×50mm) were prepared in the following manner: (1) wash with ddH_2_O; (2) wash with isopropanol; (3) wash with ddH_2_O; (4) wash with isopropanol; (5) rinse with ddH_2_O; (6) dry under a pressurised air stream. A silicone gasket with 6 cavities was placed on top/centre of the coverslip to form reaction holes. A total amount of 19 µl of working buffer 1 (20 mM HEPES pH 8, 150 mM KCl, 2 mM MgCl_2,_ 5 mM DTT) for hTFIIIC alone and working buffer 2 (20 mM HEPES pH 8, 200 mM KCl, 2 mM MgCl_2,_ 5 mM DTT) for hTFIIIC-DNA sample was applied to each reaction hole for autofocus stabilization before every measurement. 1 µl of the sample at an initial concentration of 400 nM was added to the working buffer 1 or 2. All measurements were performed using a Refeyn Two^MP^ mass photometer (Refeyn Ltd, Oxford, UK). Videos of 1 min with regular image size (150×59 binned pixels, 10.9um x 4.3um imaged area, 46.3um2 detection area) were recorded using the Acquire^MP^ software (Refeyn Ltd, version 2.4.0) and data analysis were performed using the Discover^MP^ software (Refeyn Ltd, version 2.4.0).

### Cryo-EM sample preparation

To reconstitute the hTFIIIC-DNA complex for cryo-EM, freshly purified hTFIIIC at an initial concentration of 3.17 µM was incubated with human TRR-TCT3-2 DNA (7.5 µM stock concentration) at a 1:1 molar ratio for 10 min at RT. The mixture was passed through a Zeba Spin desalting column previously equilibrated with EM Buffer (20 mM HEPES pH 8, 225 mM KCl, 2 mM MgCl_2_, 5 mM DTT). The sample was left at RT for 30 min, then mixed 0.8:0.2 (v/v) with 0.5% (w/v) of octyl-β-glucoside (previously dissolved in EM buffer) to a final concentration of 0.1% (w/v/) octyl-β-glucoside, applied to freshly plasma-cleaned 200-mesh Au R2/2 grids (UltrAuFoil) and plunge-frozen in liquid ethane using a Vitrobot Mark IV (Thermo Fisher Scientific), which was set to 100% humidity and 6 ⁰C. To prepare the cryo-EM sample of DNA-unbound hTFIIIC, the same protocol was used without using the desalting column step.

### Cryo-EM data acquisition and processing

Cryo-EM data of the hTFIIIC-DNA sample was collected on a Titan Krios G3 (Thermo Fisher Scientific) operated at 300 keV, equipped with a Gatan K3 detector and energy filter. A magnification of x105,000 corresponding to a physical 0,822 Å per pixel was used. A total amount of 11025 image stacks of 40 frames were collected in counting mode with a total electron dose of 42,8e/Å^2^ at defocus range from 0.7-1.7 µm using SerialEM^50^. The dataset of DNA-unbound hTFIIIC (11,063 micrographs) was collected with the same imaging parameters (Table S1).

Micrographs were initially preprocessed using WARP^51^. Particle coordinates were obtained by using the BoxNet2_20180918 model (without retraining) implemented in WARP. Subsequently, micrographs were preprocessed in RELION 3.1.3^52^ by using its implementation of MotionCor2^53^. CTF parameters were derived using Gctf^54^. An initial set of 530,171 particles identified by WARP was extracted with a boxsize of 480 pixels and imported into cryoSPARC 3.3.2^55^ for all subsequent classication and refinement steps. 2D classification identified two sets of particles that correspond to τA and τB-DNA subcomplexes. Each set of particles was used to train neural networks (conv127 model) for particle picking in TOPAZ^56^. For τA, a first round of particle picking in TOPAZ yielded 280,410 particles. An initial τA map was generated by using particles that showed high-resolution features in 2D classifiction. In addition, new particles in 2D classes with high resolution features were used for a second round of neural network training and particle picking in TOPAZ, which yielded 365,063 particles. Three rounds of heterogeneous refinement were performed to classify this final set of particles. The *ab initio* map of τA and three “junk maps”, previously generated in the first round of TOPAZ training, were used for the heterogeneous refinement step as input maps. A final Non-Uniform (NU) refinement applied to the best class from the last heterogeneous refinement, containing 55,079 particles, reached an overall resolution of 3.5 Å. For the τB-DNA subcomplex, a similar strategy was applied. After the first training step, 517,116 picked particles were subjected to 2D classification. The particles that belong to classes with high-resolution features were used for a subsequent TOPAZ training, while bad 2D classes were used to generate “junk maps”. A second round of TOPAZ training and picking generated 762,590 particles. A 2D classification step was applied, from which an *ab initio* τB-DNA map was generated and used for a heterogeneous refinement in the following step. 206,812 particles were further sorted by 2D classification and *ab initio* reconstruction (three classes). NU-refinement was applied to one class with 99,217 particles reaching an overall resolution of 3.3Å. To further classify these particles, the continuous reconstruction program cryoDRGN^57^ was used. The particles were binned to a box size of 256×256 px. A network with an architecture of 3 hidden layers with 512 neurons per layer for encoder and decoder and latent space size 8 was trained for 50 epochs. The resulting cluster/s averages were mapped back to derive cryo-EM density maps. Based on visual inspection of these maps a set of 35,379 particles was selected and subjected to NU refinement in cryoSPARC. In addition, a local refinement of each monomer in the τB-DNA map was performed yielding in both cases a 3.2 Å cryo-EM map. The final maps were post-processed using DeepEMhancer^58^.

### TFIIIC model building and refinement

Coot^59^ was used for building of structural models. AlphaFold^60^ predicted structures of all the human TFIIIC subunits were retrieved from the AlphaFold Protein Structure Database: TFIIIC220 (Identifier: AF-Q12789-F1), TFIIIC110 (Identifier: AF-Q8WUA4-F1), TFIIIC102 (Identifier: AF-Q9Y5Q9-F1), TFIIIC90 (Identifier: AF-Q9UKN8-F1), TFIIIC63 (Identifier: AF-Q9Y5Q8-F1), TFIIIC35 (Identifier: AF-Q969F1-F1) and fitted into the density maps using ChimeraX^61^. A B-DNA model was placed into the density with self-restraints. AlphaFold-multimer^62^ was used to assess regions with low resolution. The models were iteratively subjected to real-space refinement in PHENIX^63^ and manual adjustment in Coot. Validation of the refined model was performed using MolProbity^64^. Surface interaction area was calculated by PISA^65^. Structural homologues were retrieved from the Protein Data Bank database using the DALI 2 server^66^. Protein-DNA interactions were analysed by NUCPLOT^67^. For τA, model building was restricted to the cryo-EM density derived from the sample in which TFIIIC was reconstituted with DNA. 3D classification and refinement of τA particles from the DNA unbound sample yielded a lower resolution cryo-EM density at 3.8 Å and indicated a preferred orientation of the particles towards the air-water interface (Table S1b). Rigid body fitting of the model of τA from the DNA-bound sample into the cryo-EM density of τA from the DNA-unbound sample indicated no major conformational changes.

### Cryo-EM mapping of particle pair distances at a single-molecule level

The mapping was performed as described in Ref.^68^ with some modifications. Briefly, the particle .star files of the final RELION 3D refinement of τA and the Nu-refinement of τB-dimer subcomplexes derived from the DNA-bound sample served as input data for the particle distance pair analysis. The names and refined coordinates of the τB-dimer complex and τA were extracted from each micrograph and organized into a distance matrix of size N_τB_× N_τA_. The particles were paired with a complementing partner by determining the smallest value in the distance matrix, saving the corresponding particles, and deleting them from the matrix to avoid multiple pairings of a particle. These steps were repeated until all particles in the rows or columns are assigned. Then, a new distance matrix is calculated for the next micrograph. Thus, the algorithm iterates through all micrographs. The distances of all pairings in all micrographs were listed and plotted in a histogram (Figure S9A).

In order to estimate the number of linked and randomly paired particles, the distance of random particles was calculated. The same particle sets from the micrographs were used however, with randomized coordinates. The simulated particle distances were obtained equally and compared to the measured distances of the particles (Figure S9A).

### B-box DNA sequence conservation

To investigate B-box DNA sequence conservation across eukaryotes, we downloaded a set of eukaryotic tRNA genes from the GtRNAdb database^69^ (Release 15, http://gtrnadb2009.ucsc.edu/). For the analysis of human tRNA genes, we used the hg38 - GRCh38 dataset retrieved from the GtRNAdb 2.0 database^70^ (Release 19, http://gtrnadb.ucsc.edu/) genes. tRNA genes with a score below 70.0 were discarded for the sequence conservation analysis across eukaryotic tRNA genes, resulting in 35,829 genes. The tRNA genes were aligned using the ‘cmalign’ program within the Inferal package^71^ and the tRNA covariance model (RF00005), retrieved from the Rfam database^72^. Given that the number of input genes exceeded the maximum number of input sequences for Infernal, the data was split, aligned separately with Infernal as four separate profiles (maximum 10,000 sequences, each), and afterwards combined using the profile alignment function in MUSCLE^73^. The alignment was manually trimmed to cover only the B-box DNA motif and adjusted and trimmed to remove column gaps, for which also 15 outliers had to be removed. The final alignment for the B-box DNA motif contained 35,814 tRNA genes from 65 eukaryotic species covering three eukaryotic supergroups. Sequence logos were generated via WebLogo^74^.

### Analysis of TFIIIC220 sequence conservation

TFIIIC220 orthologs were obtained from the NCBI reference sequence database^75^ via DELTA-BLAST^76^ using human TFIIIC220 as query sequence. The retrieved protein sequence hits were filtered to reduce the bias on found opisthokont orthologs and, at the same time, cover a broad range of eukaryotic species (metazoan: 29, fungi: 29, amoebozoa: 1, plants: 59, discoba: 2). A multiple sequence alignment was generated with MAFFT^77^ and subjected to the residue sequence conservation analysis via Scorecons^78^, for which the entropic (7 types) scoring method was used.

