## Supplementary_Information for "Structural insights into human TFIIIC promoter recognition"

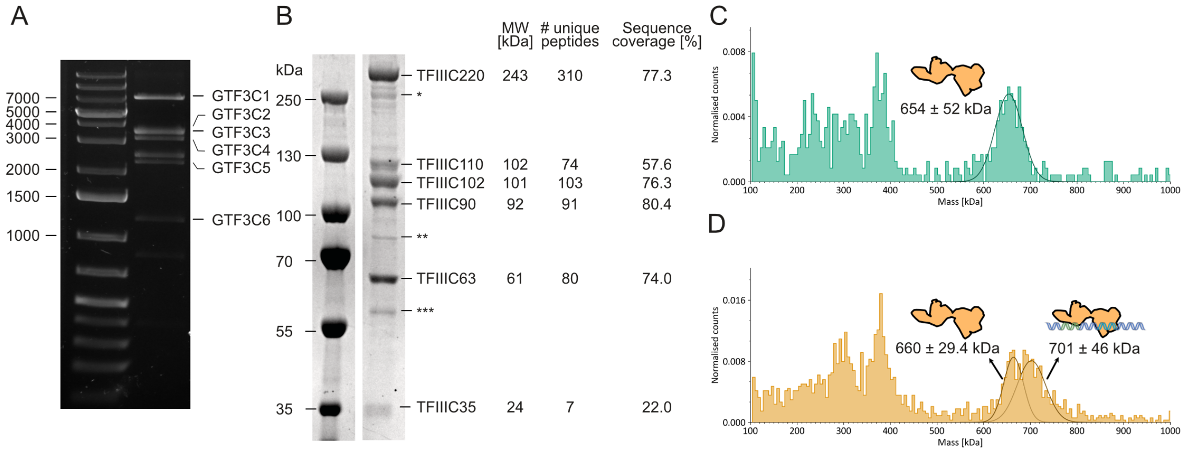

**Figure S1. Cloning, purification and mass photometry of hTFIIIC, related to Figure 1.**
**(A)** Analysis of the pBIG2ab vector containing all the hTFIIIC genes by restriction digest (SwaI) and gel electrophoresis.
**(B)** SDS-PAGE of the purified hTFIIIC complex. All the subunits were identified by mass spectrometry. (*) indicates TFIIIC220 degradation products, (**) shows the presence of 70 kDa heat shock protein from the expression host, and (***) shows α- and β-tubulins from the host.
**(C)** Mass photometry analysis of hTFIIIC. The mass distribution histogram is shown together with a schematic representation of hTFIIIC next to the peak with a molecular weight of 654 ± 52 kDa.
**(D)** Mass photometry analysis of DNA-bound hTFIIIC. Mass distribution histogram of a sample containing hTFIIIC and DNA. A schematic presentation of hTFIIIC unbound (left) and bound (right) to DNA is shown next to the peaks corresponding to 660 ± 29.4 kDa and 701 ± 46 kDa, respectively.

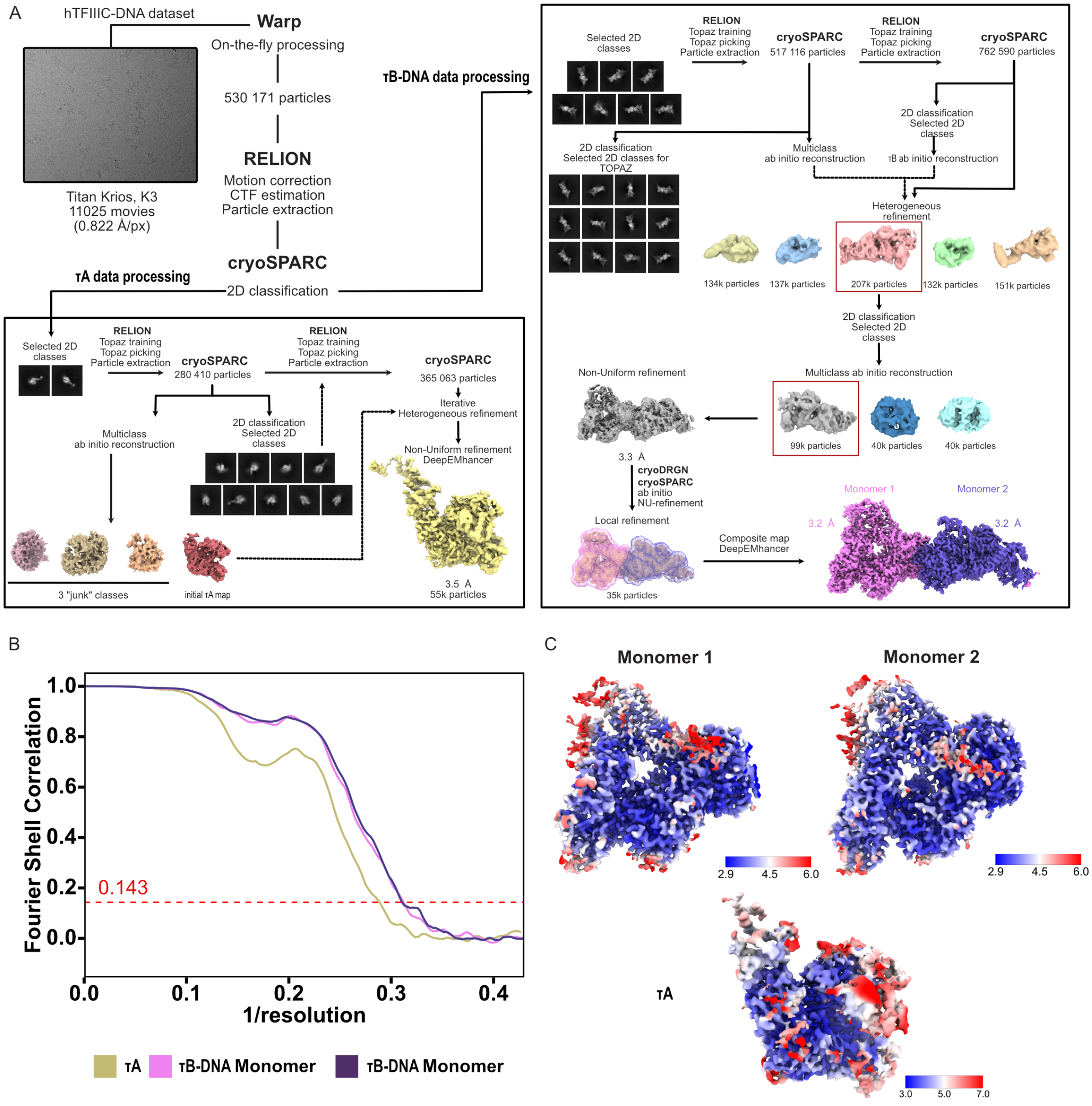

**Figure S2. Cryo-EM data processing and quality assessment of hTFIIIC, related to Figure 1, 2, 3.**

**(A)** Processing pipeline of the hTFIIIC-DNA Titan Krios dataset. A representative micrograph, 2D classes, 3D classes from heterogeneous refinement and NU-refinement maps of τA (bottom left) and τB-DNA (right) from CryoSPARC, post-processed DeepEMhancer maps are shown. **(B)** FSC curves of the τA, monomer 1 and monomer 2 from the dimeric τB-DNA map show a final resolution of 3.5 Å, 3.2 Å and 3.2 Å, respectively (FSC = 0.143). **(C)** Local resolution estimation of τA, monomer 1 and monomer 2 implemented in CryoSPARC.

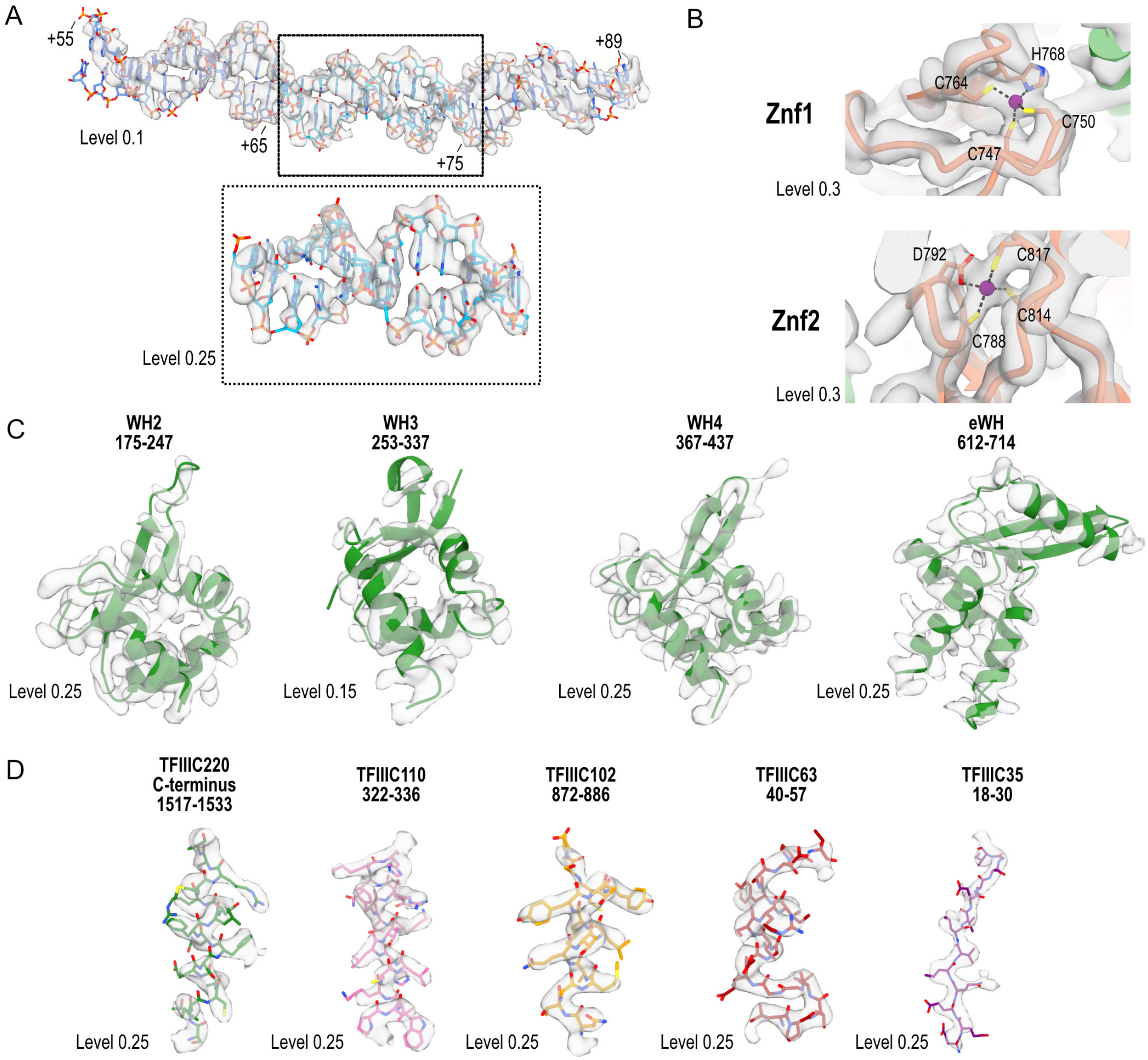

**Figure S3. Quality of map densities, related to Figure 1, 2, 3.**
Exemplary cryo-EM densities and refined models of:
**(A)** DNA, including the B box (bottom panel),
**(B)** Znf1 and Znf2,
**(C)** WH2-4 and eWH,
**(D)** TFIIIC220 C-terminus, TFIIIC110, TFIIIC102, TFIIIC90, TFIIIC63, TFIIIC35. The threshold level is indicated next to the model.

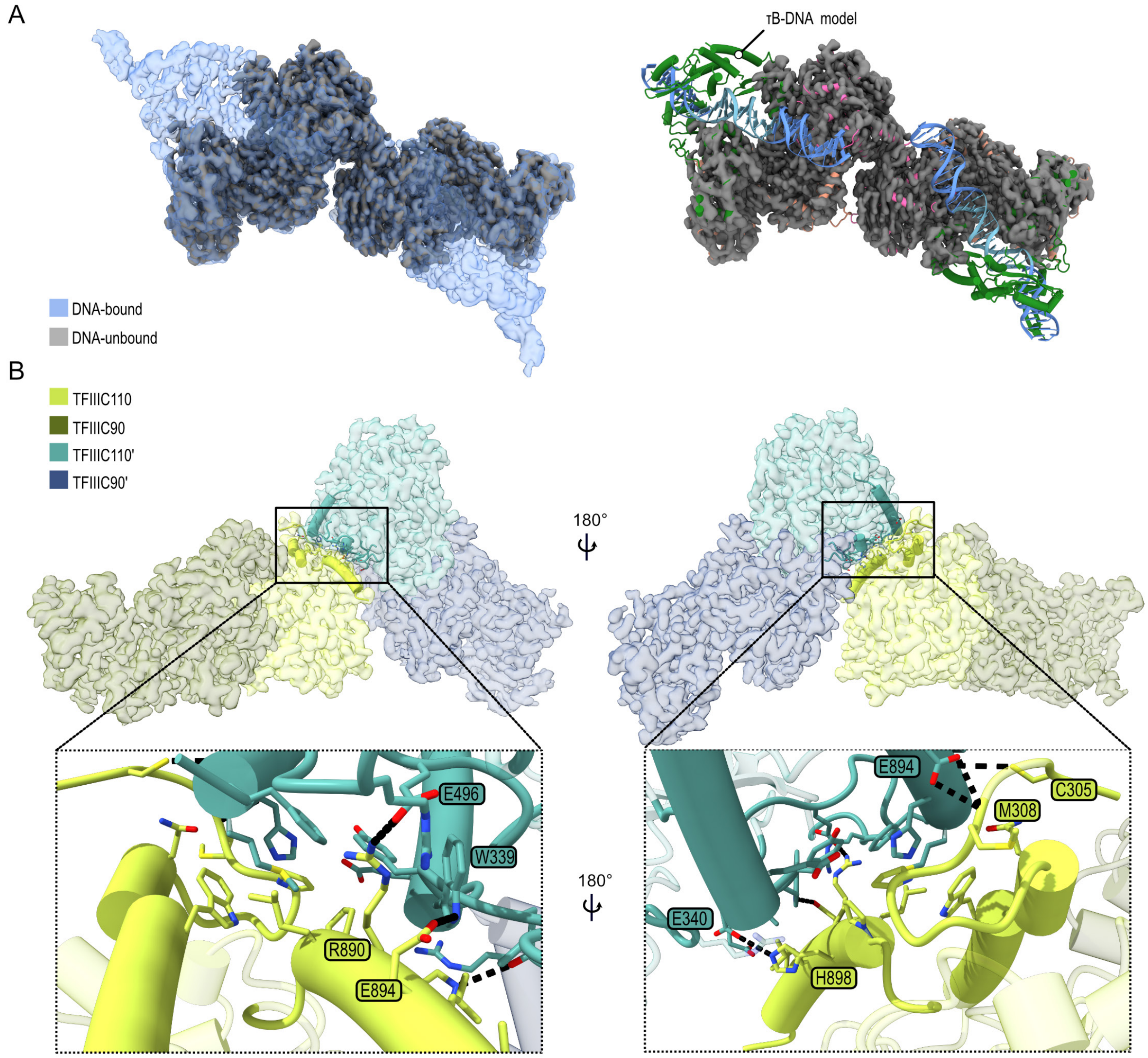

**Figure S4. Analysis of τB dimerization, related to Figure 1.**

**(A)** Left, Cryo-EM density map of DNA-unbound hTFIIIC (grey), obtained from a separate dataset without DNA, superimposed with DNA-bound TFIIIC map (blue and transparent). Right, superimposition of DNA-unbound TFIIIC map with the refined DNA-bound TFIIIC model, represented as cartoon.

**(B)** Interface interaction between τB monomers. Insets show the amino acids, which take part in the interaction. Black dashed lines represent polar interactions with the amino acids in coloured boxes. The amino acids participating in the interactions were identified by ChimeraX "interfaces" command line and PDBePISA.

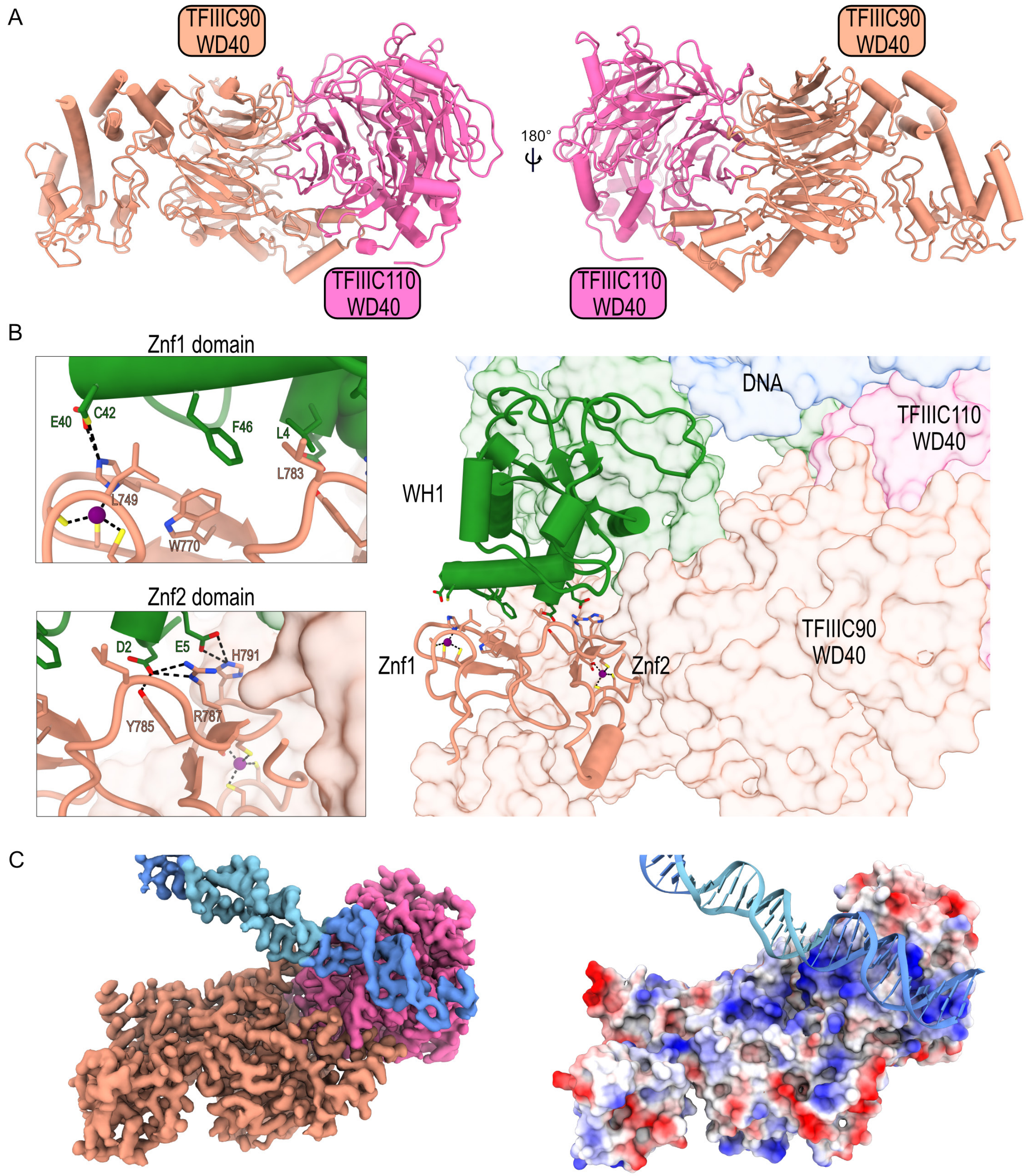

**Figure S5. Structure and DNA interaction of the human τB core, related to Figure 1, 2.**
**(A)** Structural model of the τB core formed by the interaction of two WD40 domains.
**(B)** Zinc fingers (Znf) 1 and 2 (cartoon representation) of the TFIIIC90 subunit interact with WH1 from the TFIIIC220 subunit. Insets show the chemical environment of this interaction.
**(C)** Left, DeepEMhancer maps of the τB core and DNA (B-box in light blue). Right, electrostatic (Coulomb) potential surface representation of the τB core showing a positively charged surface of the TFIIIC110-WD40 domain interacting with the DNA downstream of the B-box.

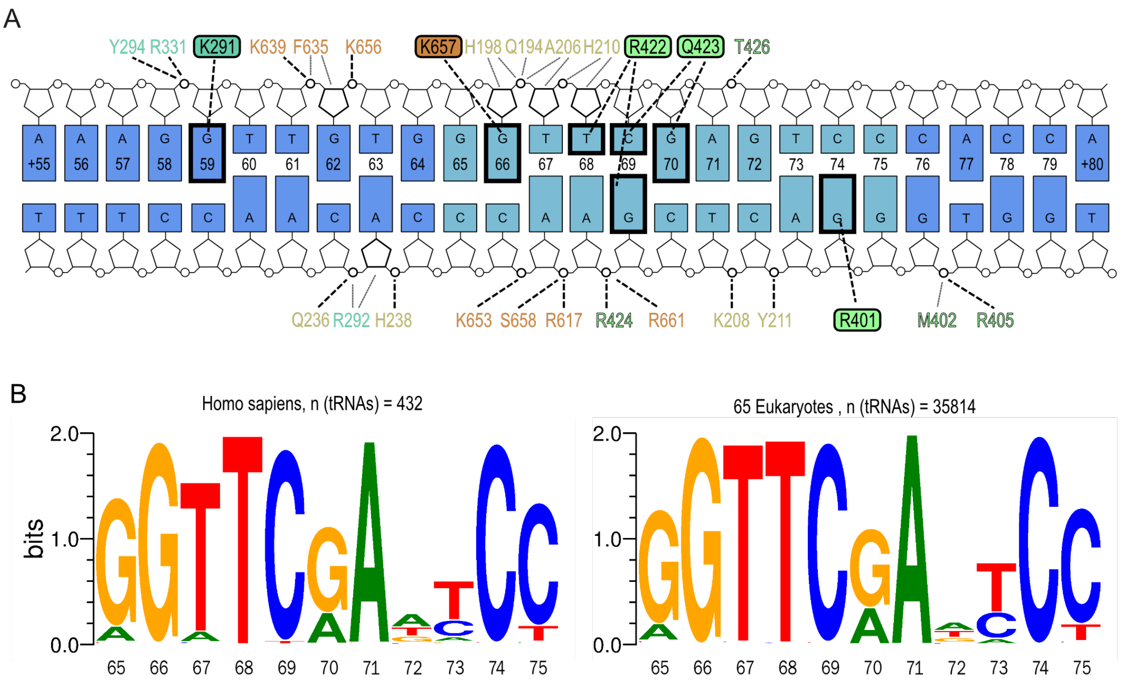

**Figure S6. DNA recognition by TFIIIC220 and analysis of B-box DNA sequence conservation, related to Figure 2.**

**(A)** Schematic of all protein-DNA interactions observed in the cryo-EM structure. Highlighted bases form H-bonds with the amino acids in coloured boxes. Domain-specific amino acid colours are the same as in Fig 2. H-bonds are depicted as bold, dashed lines and apolar contacts are shown as grey, thin lines.

**(B)** B-box DNA sequence conservation across tRNA genes from human (left panel) and 65 eukaryotic species covering metazoa (44 taxa), fungi (11), plantaea (9), and kinetoplastida (1) (right panel). The human TRR-TCT3-2 tRNA ($tRNA_{Arg}$) gene was used as a reference for the numbering of the nucleotide positions (shown below the sequence logos).

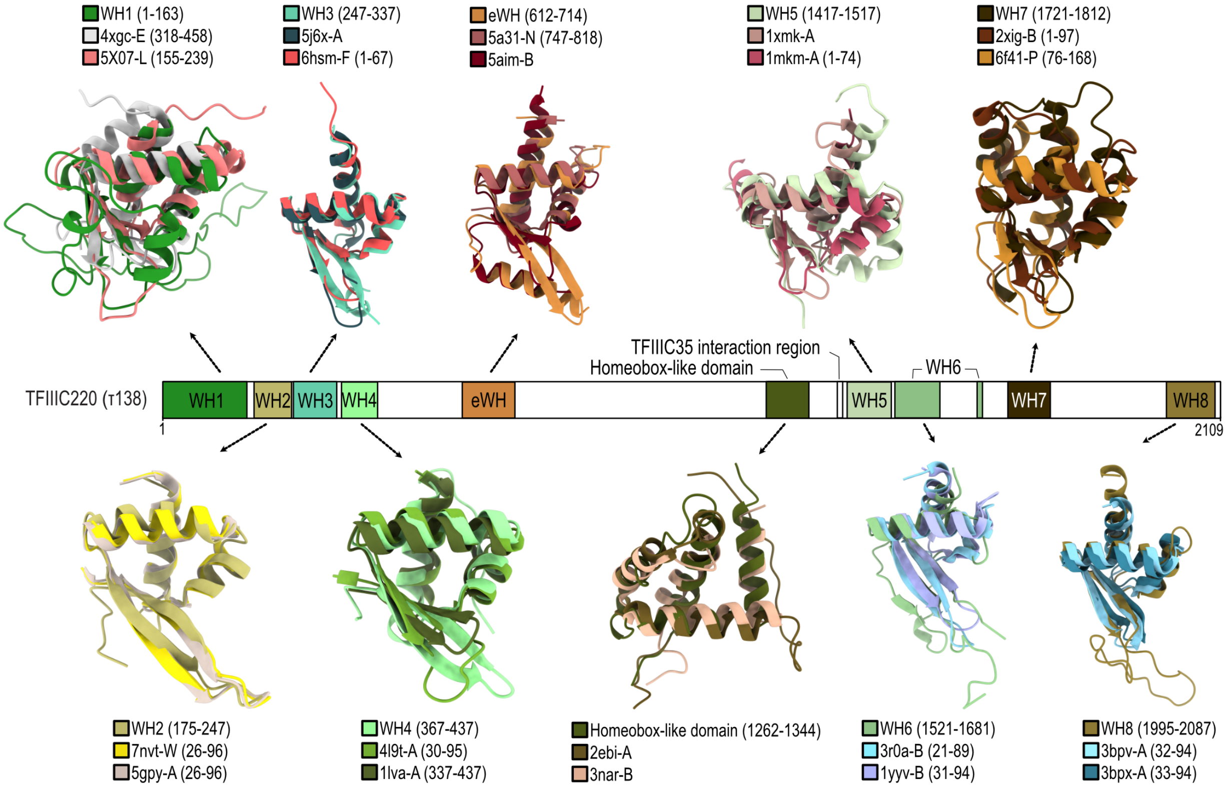

**Figure S7. TFIIIC220 subunit is mostly composed of winged-helix domains, related to Figure 2, 3.**

Superimposition of the structural models of the TFIIIC220 domains and two of their closest structural homologues retrieved from the Protein Data Bank database using the DALI server. The PDB IDs are written above (top models) and below (bottom models) each superimposition. The amino acid range for each domain is indicated in brackets. The colors of the TFIIIC220 domains are the same as in the domain architecture (middle).

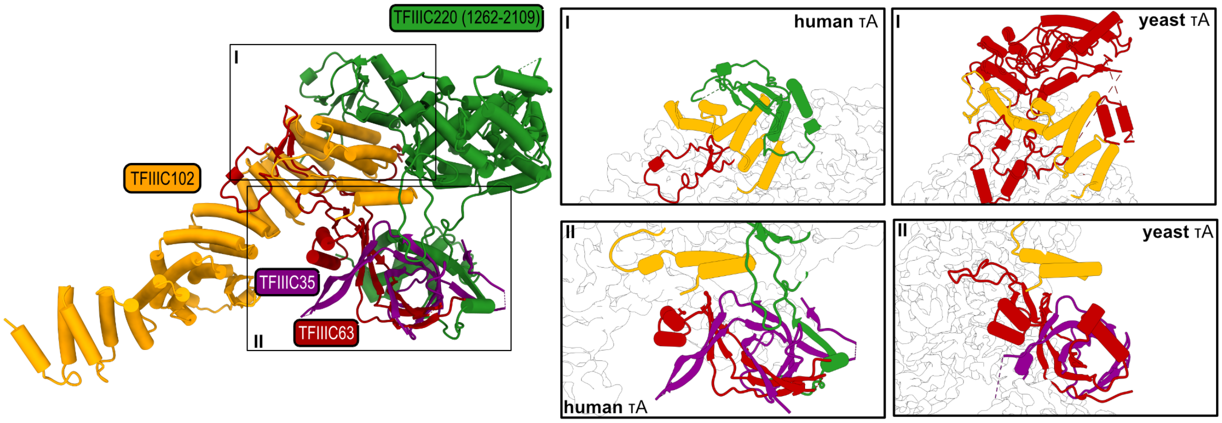

**Figure S8. Comparison between human τA and yeast τA, related to Figure 3.**
Overview of the interaction between TFIIIC102, TFIIIC63 and TFIIIC35 with the TFIIIC220 C-terminal moeity. (I) TFIIIC102 C-terminus TPR domain interacts with WH6 TFIIIC220 in human (left), while in yeast the τ131 C-terminus TPR domain interacts with the DNA-binding domain (DBD) of τ95 (right). (II) The WH8 and the TFIIIC35 IR of TFIIIC220 interact with TFIIICC63-TFIIIC35 dimer (left), absent in the yeast τA (right) (PDB ID: 6YJ6).

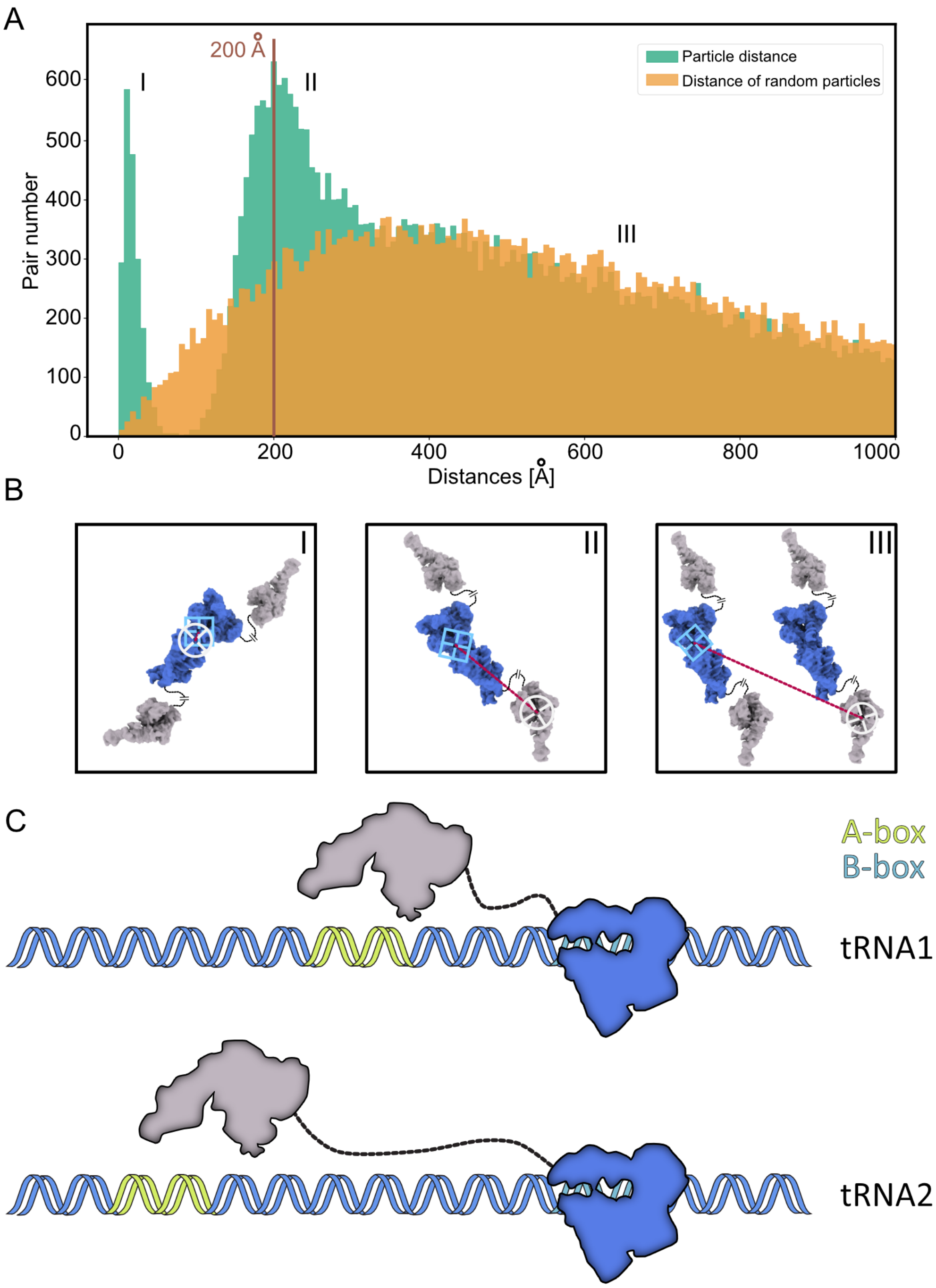

**(A)** Histogram showing the distribution of distances between τA and τB-dimer in the micrograph (particle distance) versus the distance of randomly generated particles as control**.**

**(B)** Insets showing schematics with a representative case with mislabeled particles (I), 200 Å distance between τA and τB-dimer from one TFIIIC molecule (II), from two different TFIIIC complexes (III).

**(C)** Schematics representing the flexibility of TFIIIC to bridge small and large distances between the A- and B-box present in different tRNAs. τB is depicted in blue and τA in grey.

**Table S1a. Cryo-EM data collection, refinement and validation statistics**

| | τB-DNA<br>DNA sample<br>Monomer 1 map<br>(EMDB-XXX)<br>(PDB XXXX) | τB-DNA dimer map<br>DNA sample<br>Composite map<br>(EMDB-XXX)<br>(PDB XXXX) | τA map<br>DNA sample<br>(EMDB-XXXX)<br>(PDB XXXX) | τB dimer map<br>No DNA sample<br>(EMDB-XXX)<br>(PDB XXXX) |
|---|---|---|---|---|
| **Data collection and processing** | | | | |
| Magnification | 105,000 | 105,000 | 105,000 | 105,00 |
| Voltage (kV) | 300 | 300 | 300 | 300 |
| Electron exposure (e–/Å$^2$) | 42.8 | 42.8 | 42.8 | 39.6 |
| Defocus range (μm) | 0.7-1.7 | 0.7-1.7 | 0.7-1.7 | 0.7-1.7 |
| Pixel size (Å) | 0.822 | 0.822 | 0.822 | 0.822 |
| Symmetry imposed | C1 | C1 | C1 | C1 |
| Initial particle images (no.) | 762,590 | 762,590 | 365,063 | 754,951 |
| Final particle images (no.) | 35,379 | 35,379 | 55,079 | 139,401 |
| Map resolution (Å) | 3.2 | 3.2 | 3.5 | 3.4 |
| FSC threshold | 0.143 | 0.143 | 0.143 | 0.143 |
| Map resolution range (Å) | 3.1-9.9 | n/a | 3.4-13.4 | n/a |
| **Refinement** | | | | |
| Initial model used (Alphafold structure prediction) | AF-Q12789-F1, AF-Q8WUA4-F1, AF-Q9UKN8-F1 | AF-Q12789-F1, AF-Q8WUA4-F1, AF-Q9UKN8-F1 | AF-Q9Y5Q9-F1, AF-Q9Y5Q8-F1, AF-Q969F1-F1 | AF-Q12789-F1, AF-Q8WUA4-F1, AF-Q9UKN8-F1 |
| Model resolution (Å) | 3.3 | 3.3 | 4 | 3.7 |
| FSC threshold | 0.5 | 0.5 | 0.5 | 0.5 |
| Map sharpening *B* factor (Å$^2$) | -75.9 | - | -114.5 | - |
| **Model composition** | | | | |
| Non-hydrogen atoms | 15932 | 31864 | 10800 | 25452 |
| Protein / Nucleotide residues | 1808 / 70 | 3616 / 140 | 1344 / 0 | 3178 / 0 |
| Ligands | 2x Zn | 4x Zn | - | 4x Zn |
| ***B* factors (Å$^2$)** | | | | |
| Protein (min/max/mean) | 27.9/185.9 /76.6 | 33.0/196.5 /78.9 | 67.3/184.0 /106.3 | 31.6/197.3/89.5 |
| Ligand (min/max/mean) | 111.0/132.7/121.8 | 104.6/131.9/118.2 | - | 173.7/219.7/196.7 |
| Nucleotide (min/max/mean) | 59.1/241.6/131.6 | 12.2/164.7 /93.7 | - | - |
| **R.m.s. deviations** | | | | |
| Bond lengths (Å) | 0.005 | 0.003 | 0.010 | 0.005 |
| Bond angles (°) | 0.803 | 0.712 | 1.260 | 0.875 |
| **Validation** | | | | |
| MolProbity score | 1.49 | 1.34 | 1.64 | 1.49 |
| Clashscore | 7.70 | 6.11 | 7.87 | 7.77 |
| Poor rotamers (%) | 0.63 | 0.25 | 0.42 | 0.5 |
| **Ramachandran plot** | | | | |
| Favored (%) | 97.7 | 97.9 | 96.7 | 97.7 |
| Allowed (%) | 2.3 | 2.0 | 3.26 | 2.3 |
| Disallowed (%) | 0.0 | 0.0 | 0.08 | 0.0 |

### Table S1b. Cryo-EM data collection τA without DNA

| | τA map<br>No DNA sample<br>(EMDB-XXX) |
|---|---|
| **Data collection and processing** | |
| Magnification | 105,00 |
| Voltage (kV) | 300 |
| Electron exposure (e–/Å$^2$) | 39.6 |
| Defocus range (μm) | 0.7-1.7 |
| Pixel size (Å) | 0.822 |
| Symmetry imposed | C1 |
| Initial particle images (no.) | 748,471 |
| Final particle images (no.) | 85,208 |
| Map resolution (Å) | 3.8 |
| FSC threshold | 0.143 |
| Map resolution range (Å) | n/a |
